## Supplementary figures and tables for "Transmission of SARS-CoV-2 from humans to animals and potential host adaptation"

L.v.D and F.B. are co-last authors.

**Keywords:**

SARS-CoV-2, COVID-19, human-to-animal transmission, mink, white-tailed deer, host-adaptation

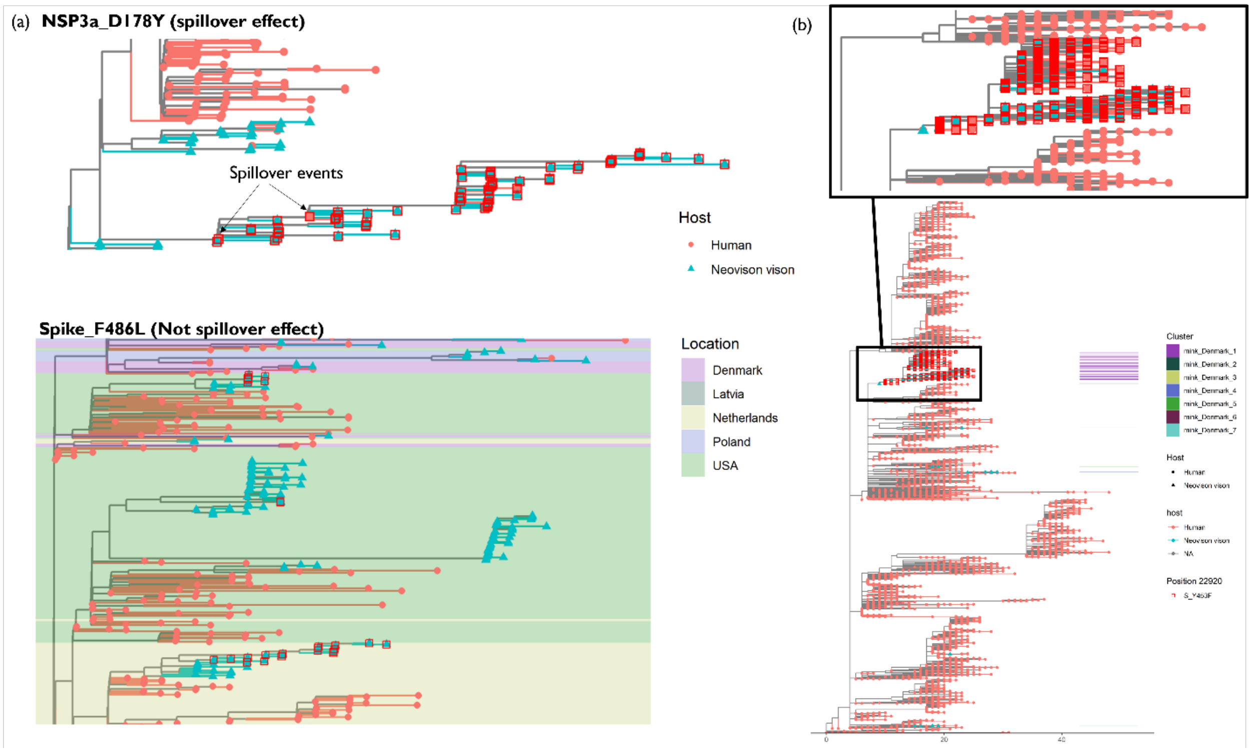


**Figure S1. Spillover effect and its lack thereof.** (a) Subtrees of mink-human phylogenies illustrating the cases where a mutation is already present in the ancestral human lineage prior to spillover (NSP3a_D178Y) and where the mutation arose in animals following spillover (Spike_F486L). (b) Danish mink-human phylogeny comprising isolates collected before 1 December 2020 (n = 15,879) and a magnified view of the mink_Denmark_1 cluster. All accessions used in this panel are provided in **Table S2j**.Tree tips and terminal branches are coloured by host of isolation. Tips where the respective mutations are present are indicated by red squares.

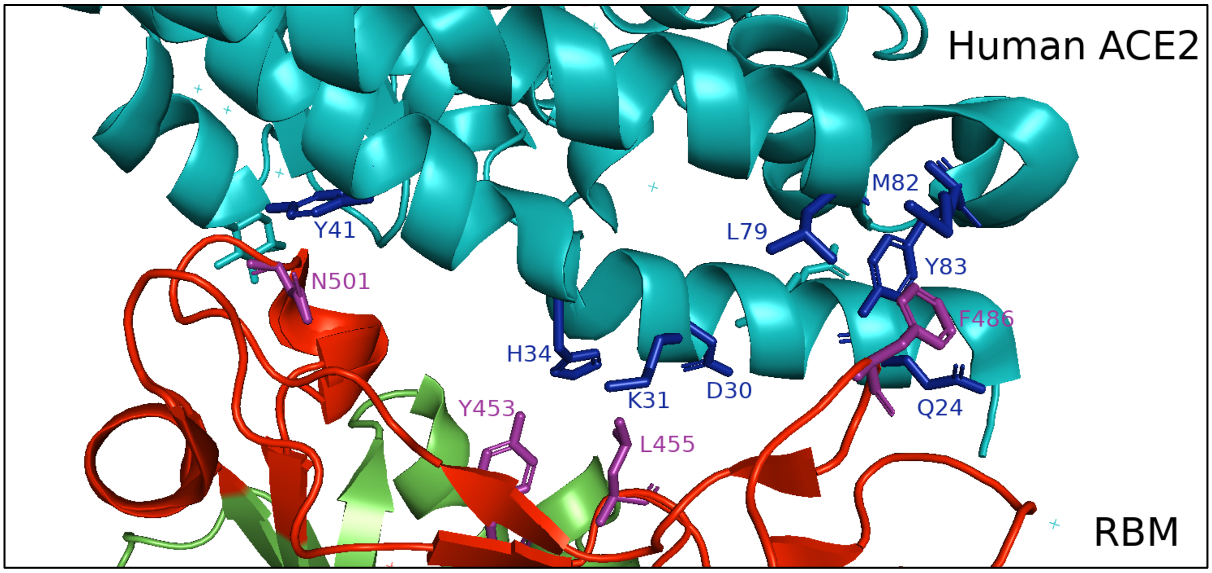


***Figure S2.*** **Protein structure of the Spike receptor binding motif (RBM) of the receptor binding domain (RBD) in complex with human ACE2**. Human ACE2, the RBM and the rest of RBD are shown in teal, red and green respectively. The wild-type amino acid residues for the candidate mutations Y453F, F486L, N501T are shown in purple. ACE2 residues reported previously to be interacting with the RBM residues L455 (in close proximity to F453), F486 and N501 are shown in blue. The Protein Data Bank (PDB) code for the SARS-CoV-2 RBM–ACE2 complex is 6M0J. This figure was rendered using *PyMOL v2.4.1*.

**Table S1. Metadata of 18 putative mink- and 31 deer-specific candidate mutations.**

**Table S2. Metadata of subsamples considered during each analysis.** (a) Metadata of 15,846 isolates representing the global context of animal-associated SARS-CoV-2 infections. (b) Metadata of 819,813 human and mink, and (c) 167,967 human and deer isolates used for manually identifying phylogenetically distinct clusters. Metadata of (d) 1,487 mink and human background 1, (e) 13,514 mink and human background 2, (f) 145 deer and human background 1, and (g) 6,670 deer and human background 2 sequences used for the allele frequency and homoplasy analyses. The metadata of the final (h) 1,068 mink and human background 1, and (i) 143 deer and human background 1 deduplicated sequences with unambiguous sampling dates used for the substitution rate analyses. (j) The 15,879 mink and human sequences from Denmark sampled prior to 1 December 2020 (see **Figure S1b**).

**Table S3. Cluster metadata for animal isolates.** These clusters were manually identified and assigned names with the format '*host-country-cluster number*'.
